# Stereotyped spatiotemporal dynamics of spontaneous activity in visual cortex prior to eye-opening

**DOI:** 10.1101/2024.06.25.600611

**Authors:** Luna Kettlewell, Audrey Sederberg, Gordon B. Smith

## Abstract

Over the course of development, functional sensory representations emerge in the visual cortex. Prior to eye-opening, modular patterns of spontaneous activity form long-range networks that may serve as a precursor for mature network organization. Although the spatial structure of these networks has been well studied, their temporal features, which may contribute to their continued plasticity and development, remain largely uncharacterized. To address this, we imaged hours of spontaneous network activity in the visual cortex of developing ferrets of both sexes utilizing a fast calcium indicator (GCaMP8m) and widefield imaging at high temporal resolution (50Hz), then segmented out spatiotemporal events. The spatial structure of this activity was highly modular, exhibiting spatially segregated active domains consistent with prior work. We found that the vast majority of events showed a clear dynamic component in which modules activated sequentially across the field of view, but only a minority of events were well-fit with a linear traveling wave. We found that spatiotemporal events occur in repeated and stereotyped motifs, reoccurring across hours of imaging. Finally, we found that the most frequently occurring single-frame spatial activity patterns were predictive of future activity patterns over hundreds of milliseconds. Together, our results demonstrate that spontaneous activity in the early developing cortex exhibits a rich spatiotemporal structure, suggesting a potential role in the maturation and refinement of future functional representations.

**Significance statement:** Understanding the temporal dynamics underlying the network structure in early development is critical for understanding network function and plasticity. By imaging hours of spontaneous cortical activity, we show strong evidence that the vast majority of spontaneous neural activity is dynamic with repeated and complex spatiotemporal patterns with stereotyped structure across hours. This suggests the potential for Hebbian learning in the development and refinement of functional visual representations. We also find that frequently occurring spatial activity patterns are predictive of subsequent activity for up to one second, which may indicate attractor dynamics in spontaneous activity. Our findings characterize key features of the temporal structure of spontaneous activity in visual cortex early in development and deepen our understanding of developing neural networks.

## Introduction

During development, functional neural networks emerge in the visual cortex that exhibit selectivity for features of the visual environment. In animals such as humans, other primates, and carnivores, these networks are organized into modular maps in which nearby neurons have similar selectivity and groups of similarly tuned modules are distributed across the visual cortex (Hubel and Wiesel 1997). Such modular maps have been identified for visual features such as edge orientation (Blasdel and Salama 1986) and direction of motion (Weliky, Bosking, and Fitzpatrick 1996), amongst many others (Issa, Trepel, and Stryker 2000; Kara and Boyd 2009; Smith, Whitney, and Fitzpatrick 2015). In the ferret, selectivity for orientation and direction emerge over the course of development, with orientation preference maps first appearing around the time of eye opening (approximately postnatal day 30-32) (Chapman, Stryker, and Bonhoeffer 1996), and direction emerging over the subsequent week (Li, Fitzpatrick, and White 2006).

Notably, the distributed and modular organization that is a hallmark of these functional maps is present in ongoing spontaneous activity at least ten days prior to eye opening (Smith et al. 2018). These early modular patterns of millimeter-scale spontaneous activity predict features of the future orientation map, but also undergo considerable refinement prior to eye opening (Smith et al. 2018). A central feature of this early spontaneous activity is the presence of long-range millimeter-scale correlations between modules.

Computational and experimental results have suggested that prior to the developmental emergence of long-range horizontal axons (Durack and Katz 1996), such long-range correlated networks can arise through purely local interactions. Models in the form of local excitation / lateral inhibition (LELI) can account for both the presence of modular functional activity as well as millimeter-scale correlations (Smith et al. 2018), and are consistent with the ability of the early cortex to self-organize unstructured inputs (Mulholland, Kaschube, and Smith 2024).

However, this prior work has focused on the spatial structure of activity patterns, leaving unexplored the temporal dynamics of early spontaneous activity, which likely have broad implications for developmental plasticity in the cortex. Various forms of spatiotemporal organization have been observed in the visual cortex in older animals or at different scales (Muller et al. 2018; 2014; Sato, Nauhaus, and Carandini 2012; Benucci, Frazor, and Carandini 2007; Siegel et al. 2012; Huang et al. 2010; Smith et al. 2015), but the spatiotemporal structure of the millimeter-scale modular spontaneous activity seen in early development remains unknown. Here we address this by recording hours of spontaneous activity in the ferret visual cortex in animals 4-9 days before eye-opening. Implementing high-speed widefield calcium imaging and a deconvolution-based analysis, we show that non-stationary temporal dynamics underlie the vast majority of large-scale spontaneous events. Our results show that the majority of events are poorly described by a linear traveling wave, but instead exhibit more complex and non-linear spatiotemporal dynamics. These dynamics showed stereotyped sets of activity patterns that re-occurred across hours of spontaneous activity. Within events, activity patterns converged to a relatively low number of possible spatial motifs, and then followed separable trajectories for many hundreds of milliseconds. Together, these results reveal for the first time the complex spatiotemporal patterns of spontaneous activity in the early developing visual cortex, revealing stereotyped sequences of activity that may suggest the presence of attractor-like dynamics.

## Results

### Early cortical activity exhibits fast temporal dynamics

In order to examine the spatiotemporal structure of millimeter-scale modular spontaneous activity in early development, we injected the neonatal ferret cortex with AAV expressing GCaMP8m between ages P10-14. We then recorded hours of spontaneous activity between ages P23-28, approximately 4-9 days before normal eye opening. To capture the temporal dynamics of spontaneous activity, we paired GCaMP8m, with its relatively fast decay kinetics, with mesoscopic imaging at 50 Hz, capturing approximately a 6 mm^2^ field of view. Experiments were performed under light isoflurane anesthesia, which in prior work preserves the millimeter-scale spatial structure of spontaneous activity seen in the awake cortex (Smith et al. 2018). For our purposes, this imaging regime also has the advantage of producing spontaneous events that tend to be separated by periods of relatively low activity, allowing us to more easily separate events in order to focus on the fast temporal dynamics within events. Consistent with prior work, we observed frequent spontaneous events (Fig. 1A) that exhibited modular and distributed millimeter-scale correlations across the field of view (FOV) (Fig. 1B). In order to better resolve the temporal structure within these events, we performed a prior-frame subtraction (PFS) deconvolution method (Stern, Shea-Brown, and Witten 2020), in which prior frame fluorescence data scaled by the decay time constant for GCaMP8m (Zhang et al. 2021) was subtracted for each imaging frame (Fig. 1C,D). Spontaneous events were then segmented into discrete periods of contiguous activity (see Methods).

**Figure 1.**
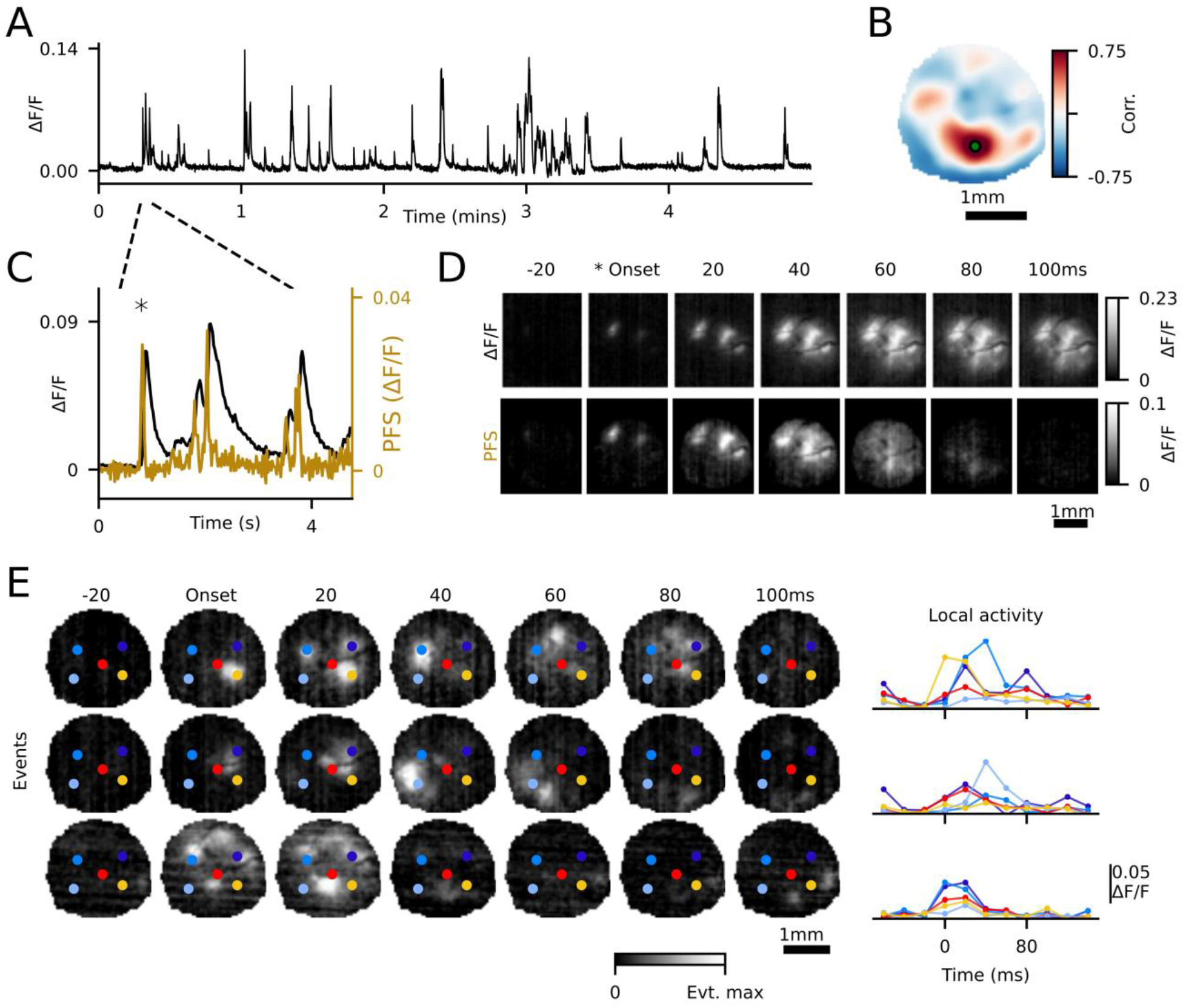
Early cortical activity exhibits fast temporal dynamics. (**A**) Spontaneous activity, averaged over field of view. (**B**) Correlation across field of view with respect to the green seed point. (**C**) Averaged activity, zoomed in from panel **A**. Left and right y-axes, respectively, show ΔF/F_0_ and a “previous frame subtraction” (PFS) deconvolution approach. (**D**) Top row shows ΔF/F_0_ and bottom row shows deconvolved activity. (**E**) Three segmented events bounded on either side by inactive frames. Each event is normalized to its maximum value. (right) averaged traces of activity within color-coded circles, highlighting dynamic activity over time.

Consistent with prior observations of spatially distributed, modular activation patterns in the developing cortex (Smith et al. 2018), we found that many events exhibited spatially segregated regions (modules) of activation (Fig. 1D,E). Notably, the spatial patterns of activated modules often shifted over tens of milliseconds within spontaneous events, revealing fast spatiotemporal dynamics. These dynamic activity patterns largely exhibited modular structure even at higher temporal resolution. We often observed that activity initiated in one module, then shifted to another module in a different location on the next imaging frame (separated by 20 ms) (Fig. 1E, top two rows). In other cases, large patterns of active modules appeared nearly simultaneously, with the same spatial pattern maintained over the course of the event (Fig. 1E, bottom). These results indicate that the improved temporal resolution of our data allows us to investigate the dynamic spatiotemporal patterns of spontaneous activity in the developing cortex.

### Spontaneous activity frequently exhibits dynamic propagation across the cortical surface

To systematically examine the spatiotemporal properties of the events, we first quantified the degree of spatial propagation within each event. We defined the *propagation area* (PA) to be the area of active pixels in our imaging FOV extending beyond the activity from the first (onset) frame of the event (see Methods, Fig. 2A,B). A large majority of events showed extensive propagation, with new activity beyond onset patterns frequently extending over 1 mm^2^ (median=0.80 mm^2^, 25th-75th=0.335-1.399 mm^2^ for 8093 events from 6 animals; Fig. 2C).

**Figure 2.**
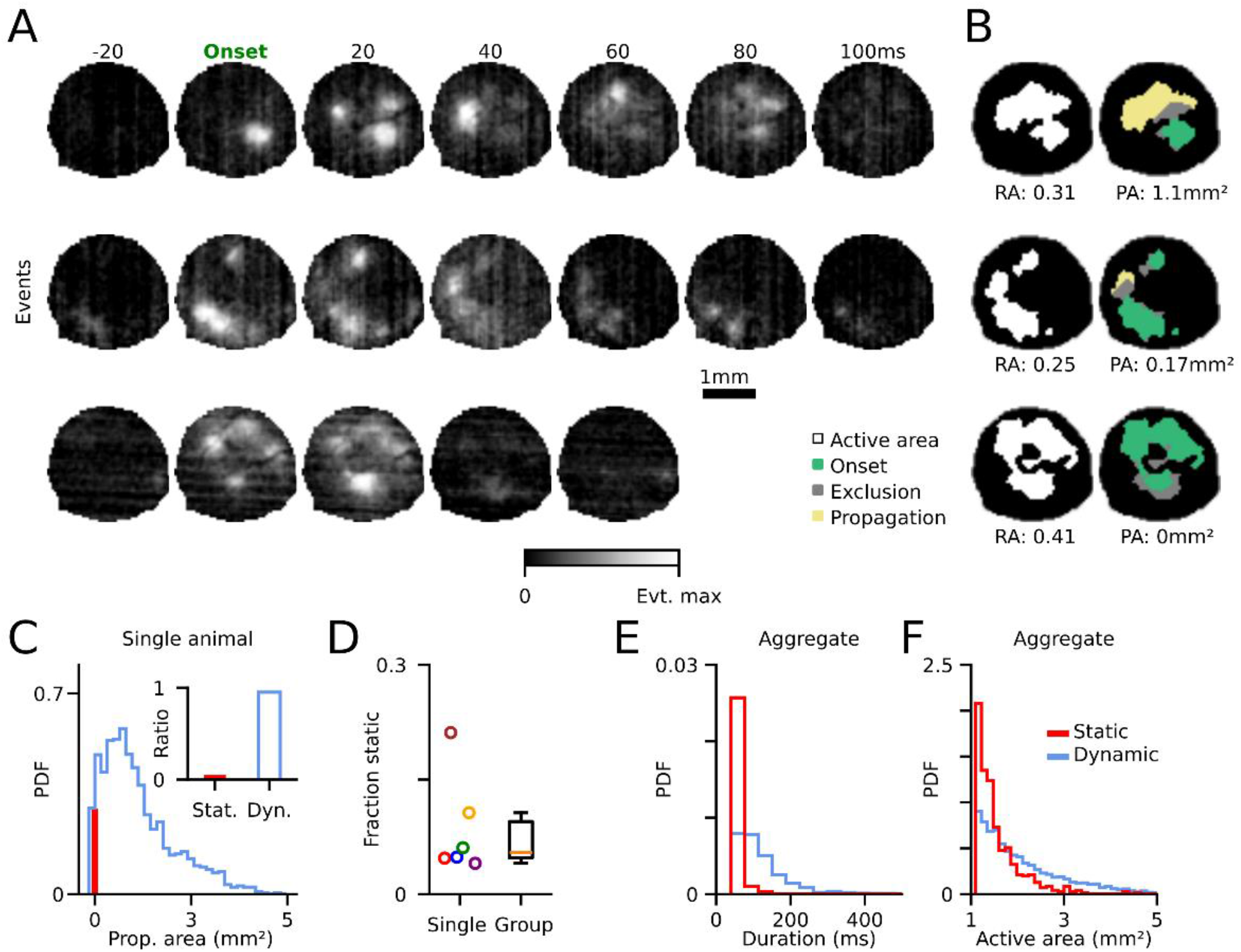
Spontaneous events exhibit propagating activity patterns across the cortex. (**A**) Three example events with varying amounts of propagation. (**B**) (left) Ratio of all active pixels during entire event (RA). (right) Area of propagating activity (PA, yellow) extending beyond onset activation (green) (see Methods). (**C**) Propagation area across all events. Red bar indicates static events (PA=0). Inset shows percentage of static and dynamic events (PA>0) for one animal. (**D**) Summary of static event percentages across animals (mean=8.6%, CI=4.3-18.5%, n=6) (example animal is blue). (**E**) Static events exhibit shorter durations than dynamic events. (**F**) Static events exhibit smaller active area than dynamic events.

To estimate the fraction of events that showed within-event spatial propagation, we categorized all events with PA > 0 as ‘dynamic’ and propagating, and all events with PA = 0 as ‘static’ and non-propagating. In static events, all activity remained localized to regions present in the initial onset of activity. Across all animals, we found that the majority of events were dynamic and only a small subset was static (mean=8.6%, range=4-21%, n=6 animals) (Fig. 2D). This indicates that dynamic events are the prevalent mode of cortical activity at this stage of development. The high frequency of dynamic propagation we observe in our data also argues that our imaging resolution is sufficient to capture the underlying temporal structure of spontaneous activity, as propagation faster than our imaging speed would appear as ‘static’, which we rarely observe. We next sought to quantify the temporal duration and total active area of these spontaneous events. Overall, static events were significantly shorter in duration than dynamic events (dynamic: median=100ms, median range=60-120ms; static: median=40ms, median range=40-40ms; n=6 animals, Mann-Whitney U test, p=9.8x10^-4^, Fig. 2E), and engaged less cortical area (dynamic: median=1.8 mm^2^, median range=1.6-2.0 mm^2^; static: median=1.4 mm^2^, IQR=1.3-1.4 mm^2^; Mann-Whitney U test, p=2.1x10^-3^, n= 6 animals, Fig. 2F). These results indicate that spontaneous events often initiate in a localized spatial region and then cascade into new regions within the cortical network. Furthermore, events rarely emerge simultaneously across a large spatial area, but rather expand in footprint dynamically across time.

### Linear traveling waves describe a minority of propagating spontaneous events

We next examined if this propagating activity moves across the cortical surface in a traveling wave, as has been observed in prior studies (Muller et al. 2014; Sato, Nauhaus, and Carandini 2012; Nauhaus et al. 2012; 2009; Grinvald et al. 1994; Benucci, Frazor, and Carandini 2007). We identified active pixels on each frame of an event and then attempted to fit their activation time with a linear traveling wave moving across the imaging FOV. To assess the quality of these linear fits, we computed a “waviness index” as done in (Siegel et al, 2012) which compares the fitting error of an event to the fitting errors of frame-permuted events, thereby maintaining spatial characteristics but destroying temporal relationships. We found that across animals, only a minority of events were well described by a propagating linear wave (median 32%, range 22-53%, n=6 animals) (Fig. 3A,B,C).

**Figure 3.**
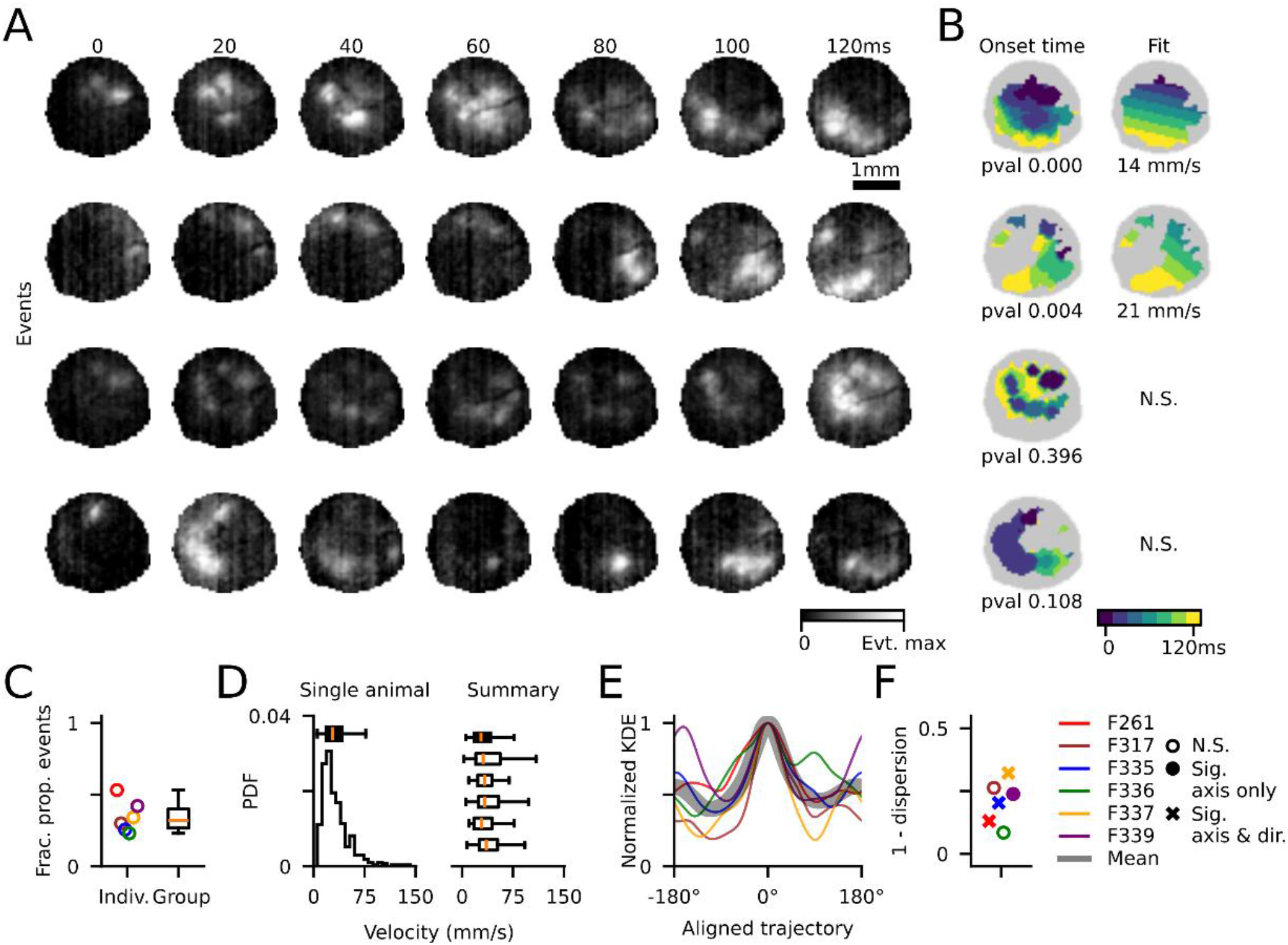
A subset of events is well described by a linear traveling wave. (**A**) Four example events. (**B**) (left) Onset time for all active pixels in an event and (right) the linear fit. Bottom two events were not well fit by a linear wave. (**C**) Fraction of events exhibiting significantly linear propagation (median 32%, range 22-53%, n=6 animals) (**D**) Wave propagation velocity estimates for linear waves. (left) Example animal and (right) all animals. (**E**) Summary of trajectories across all animals, aligned to most common propagation direction and normalized within animal. Black shaded line is the mean across animals. (**F**) 1 - Angular dispersion of wave propagation axis for linear waves in each animal. Distributions were tested for significant axis of propagation (180° space) and if they had a preferred trajectory along the axis. Statistically significant axis of propagation with no preferred direction is shown with a filled circle (p<0.05, Rayleigh test), and axes with preferred trajectory are marked with an “x” (p<0.05, binomial test).

Of these events that were significantly fit by a linear wave, the propagation velocity was fairly consistent across animals, with a median velocity of 32 mm/s (5-95th=12-114 mm/s, 1432 events across 6 animals, Fig. 3D). Linearly propagating events traveled across a range of directions within each animal. In most animals, however, there was a weak but significant bias in preferred direction (Fig. 3E, Supplemental Fig. 1). Four of six imaged animals exhibited a significant axial bias (180⁰) in linear wave propagation direction (p<0.05 Rayleigh test, see Supplemental table 2), three of which also exhibited a significant directional bias (post-hoc binomial test, p<0.05) (Fig. 3F). Thus, despite the high fraction of spontaneous events that exhibit fast propagation on the scale of hundreds of milliseconds, relatively few of these events exhibit simple linear dynamics. Rather, the majority of early spontaneous events show more complex and non-linear spatiotemporal trajectories during early development.

### Spontaneous activity shows stereotyped spatiotemporal patterns that are conserved across hours

In order to better understand the complex spatiotemporal dynamics of spontaneous activity in the early developing cortex, we next examined if similar patterns of activation occurred across multiple events. To achieve this, we computed inter-event correlations by combining all active frames within each event, allowing us to capture the overall similarity in activation patterns while incorporating both spatial and temporal components (see Methods). To illustrate this approach, we selected two sets of events that showed high event-level correlations within sets and low correlations between sets (Fig. 4A) and computed the spatial frame-to-frame correlations for all frames across these events (Fig. 4B). This revealed high correlations between aligned frames across correlated events, indicating that the spatial pattern of activity followed a highly similar progression across time, which was repeated in each instance of the event.

**Figure 4.**
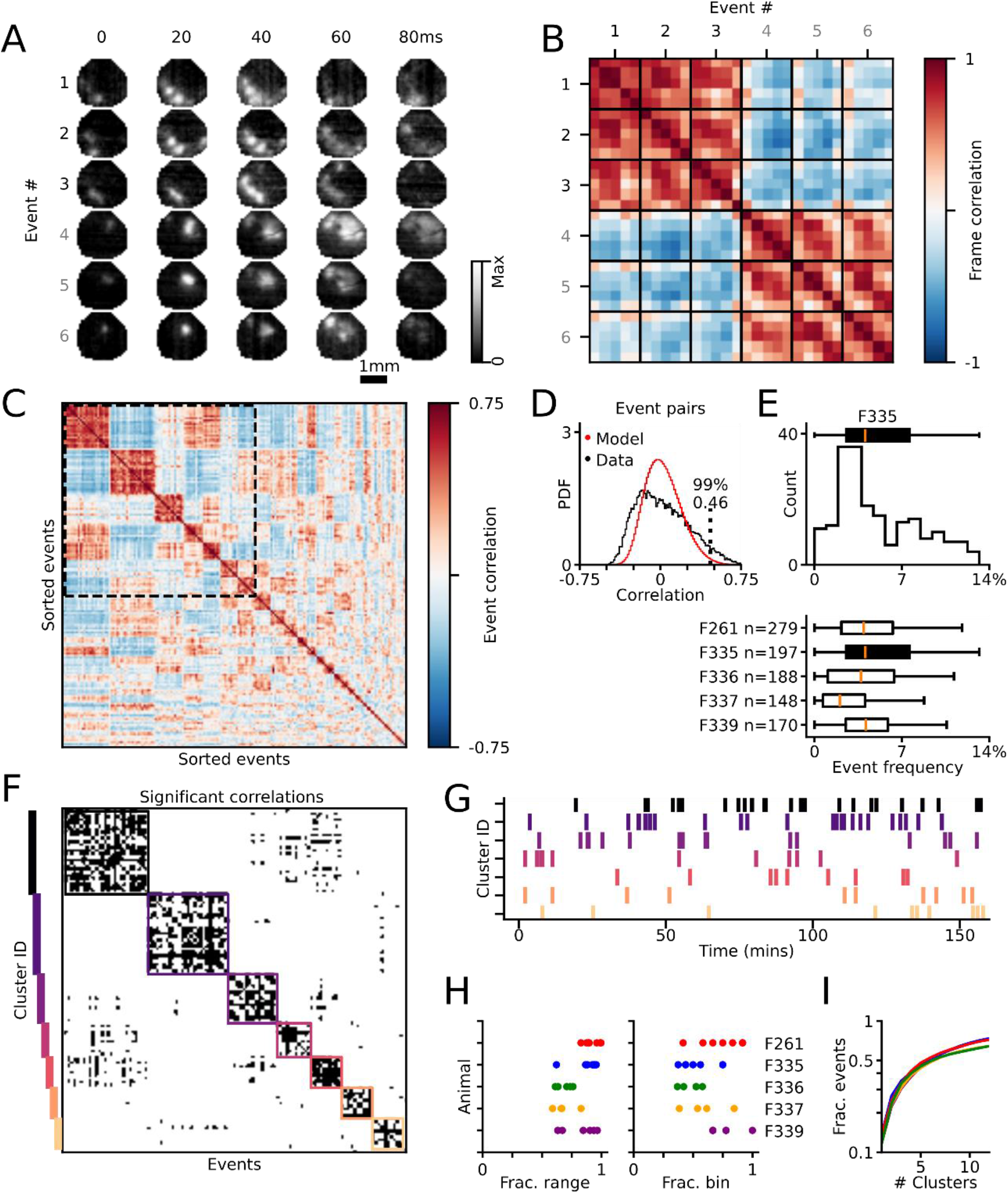
Spatiotemporal patterns within events are persistent across hours. (**A**) Two example event clusters show two distinct spatiotemporal patterns. (**B**) Frame to frame correlations of events in **A**. There is high correlation within clusters and negative correlation between clusters. Additionally, there is higher correlation along diagonals, reflective of shared dynamic activity across time. (**C**) Clustered correlation matrix of all 100ms events in example animal. (**D**) Distribution of event-event correlations for actual (black) and random model (red, see Methods). Threshold for significant correlations (99^th^ percentile of model) indicated with dashed line. (**E**) (top) Distribution of event repeat frequency for all 100ms events, for example animal shown in A-D and (bottom) for all animals. (**F**) Thresholded and zoomed-in version of correlation matrix in **C**. Color-coded boxes drawn around identified clusters. (**G**) Initiation times of events found in each cluster. (**H**) (left) Fraction of recording time spanned by instance of repeated sequence motifs. (right) Fraction of ten-minute bins with at least one instance of a repeated event. (**I**) Fraction of events included as a function of increasing cluster number.

To extend this observation to all events, we used a clustering approach to identify groups of similar spatiotemporal activity patterns across events. Specifically, we computed the event-event correlation matrix across all events with duration of five frames (100 ms) and identified clusters of highly correlated events using a greedy algorithm (see Methods), which revealed subsets of spatiotemporally distinct event patterns (Fig. 4C). We then thresholded these correlations using the 99th percentile of a control shuffle distribution to identify significant correlations (Fig. 4D). Using this, we then determined the frequency with which each event was repeated, taken as the percentage of all events that were above this threshold. We found that events had a median frequency of 3.6% (25-75th=1.8-6.1%, n=982 events across 5 animals, Fig. 4E), indicating that most events occurred as repeated instances of similar spatiotemporal patterns.

We next examined the temporal distribution of repeated instances of similar spatiotemporal event patterns across the hours of imaging in our dataset. Using the most frequently observed clusters of similarly structured events (Fig. 4F), we observed that instances of each cluster occurred throughout the majority of the imaging session (Fig. 4G, see Supplemental Fig. 2). Across animals, significantly similar events were observed spanning hours of imaging, covering the majority of time within an imaging session (percentage imaging time spanned by event instances: median=89%, 25-75th=78-95%, n=30 clusters across 5 animals, Fig. 4H, *left*), and occurring within the majority of 10-minute segments of binned spontaneous activity (percentage segments with identified event: median=67%, 25-75th=55-80%, n=30 clusters across 5 animals, Fig. 4H, *right*). Notably, the spatiotemporal similarity across similar event motifs identified through this clustering approach explained a majority of the data, with over half of all 100 ms events falling within 8 identified clusters (Fig. 4I, range 56- 61%, n=5 animals). Collectively, these analyses demonstrate that spontaneous activity in the developing cortex contains distinct motifs of spatiotemporal activity patterns, which occur repeatedly across hours of cortical activity.

### Spontaneous activity motifs are predictive of future activity patterns over hundreds of milliseconds

One of the most prominent features of early spontaneous activity is the presence of long-range correlations, indicating that certain spatial modules tend to be co-active (Smith et al. 2018) (Fig. 1B). This is supported on a finer timescale by our finding of repeated spatiotemporal motifs in spontaneous activity. This raises the possibility that activity within an event propagates to a limited subset of distinct spatiotemporal trajectories, namely, to modules within the current network. Alternatively, the trajectories of spontaneous events could intermingle, and any given pattern of activity within an event could lead into a broad range of potential subsequent patterns. We therefore sought to investigate the degree to which the activity at a single time point could provide information about the structure of activity at both preceding and subsequent times.

We reasoned that by identifying highly repeated single frame activity patterns, we would be able to determine if spontaneous events follow a consistent trajectory through activity states, or if conserved single frame activity patterns could be engaged from many different trajectories. To this end, we first identified for each animal single-frame template motifs based on the seven most common single frame activity patterns (Fig. 5A; see Methods). We then sorted all events into groups based on their single-frame similarity to these template motifs, with the group label determined by the best matching template (see Methods). Comparing frame-to-frame correlations both within and across groups revealed a clear separation in activity patterns, as expected (Fig. 5B).

**Figure 5.**
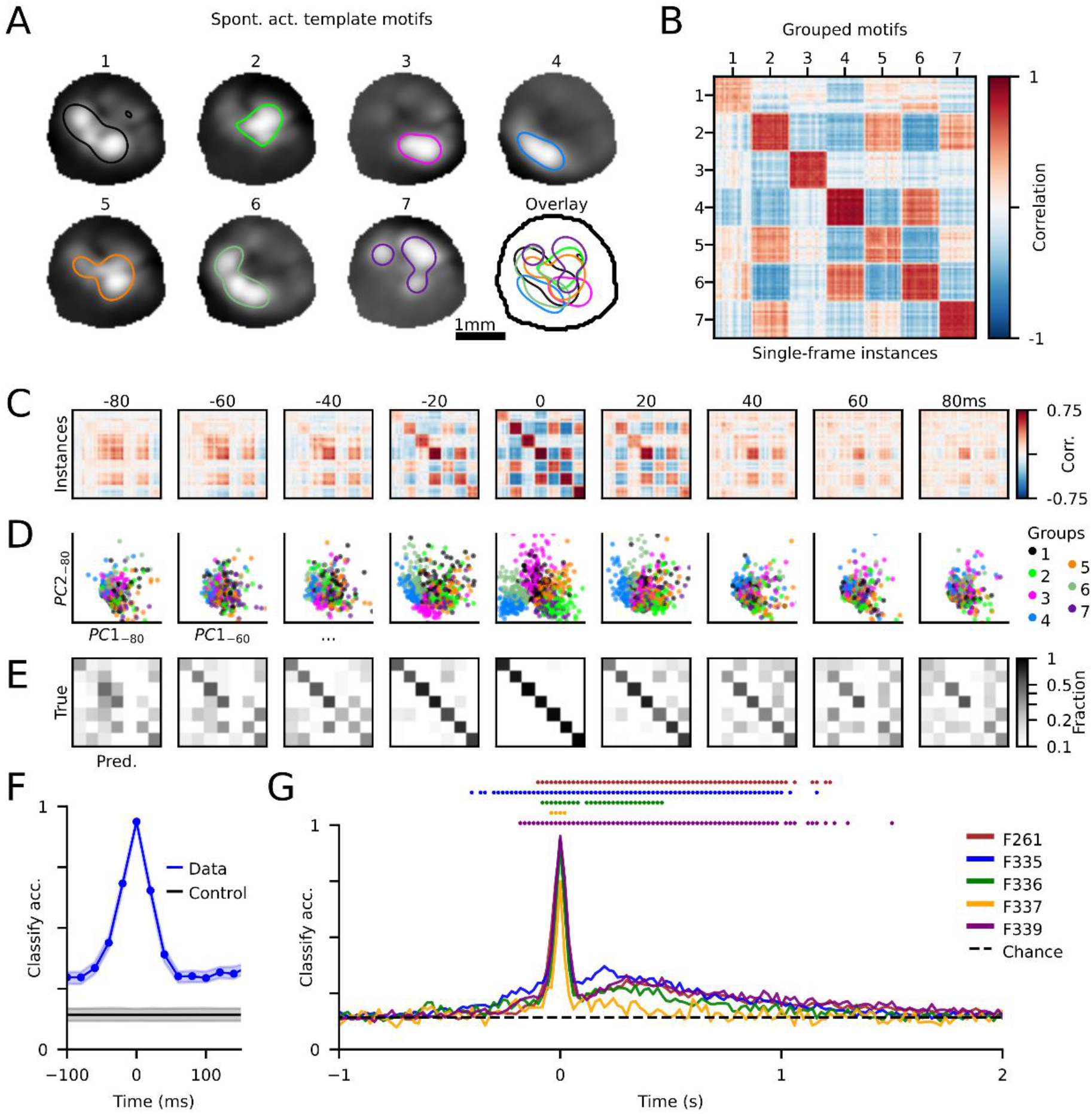
Spontaneous activity motifs are predictive of future activity patterns over hundreds of milliseconds. (**A**) Template frame motifs, showing the average single-frame group activity, identified by finding highly repeated instances in single frames of activity. Contours are drawn to illustrate each template’s active area. (**B**) Single-frame correlation matrix, sorted by group. (**C**) Single-frame correlation matrices for time points −80ms to +80ms, from template groups sorted as in **B**. (**D**) Group separability shown for all time points, plotted as first two principle components of each time point. (**E**) Group classifier performance from −80ms to +80ms. (**F**) Classification accuracy across time for example animal shown in **A-E** and control scores’ mean performance of shuffled groups (CI=5-95%). (**G**) Classification accuracy for all animals. Dots indicate time-points for each animal significantly above shuffled control accuracy (p<0.05).

In order to understand how these patterns of activity change over time, we examined the frame-to-frame correlations for each sequence at each time steps up to ±80 ms away from the template frame using the group labels defined at the template frame (t=0). The strong organization in the correlation matrices when sorted by group label appeared to retain some structure even ±80 ms away, as evidenced by the prevalence of positive correlations near the diagonal (Fig. 5C). To better understand how groups remained separable across time, we projected activity patterns into the principal component space computed at each time step. Single events are plotted by their coordinates in the first two PCs and colored by the group label at t=0 (Fig. 5D). Consistent with the strong organization evident in the sorted correlation tables, projections into PC space were clearly separable at t=0 and at each time step ±80 ms away, with increasing across-group overlap becoming apparent at more distant time points (Fig. 5D).

To quantify this separability between activity trajectories and to determine the degree to which they diverge over time, we trained a classifier to predict the group label (defined at t=0) at each time step ±2000 ms away, using the first eight principal components of events as the input (see Methods). Confusion matrices, which report the five-fold cross-validation test accuracy, revealed that although classifier performance drops off rapidly with time, it remains above chance levels for at least many tens of milliseconds within an event (Fig. 5E), which was confirmed by comparing classifier accuracy to shuffled control data (Fig. 5F). Across most animals, we found that statistically significant classifier performance extended well beyond the template frame, including up to one second later (Fig. 5G). Notably, although the classifier was able to predict group identity for frames preceding the template in all cases, classifier performance exhibited a pronounced asymmetry with respect to time and had significant accuracy extending much further in time following the template. This suggests that after passing through an activity state highly similar to a template—which were identified as the most common single-frame activity patterns—the space of trajectories of spontaneous activity becomes narrower and more conserved, thereby leading to significant classifier performance over long timescales.

Collectively, these results indicate that the complex and largely non-linear patterns of propagating activity within spontaneous events are not random, but instead tend to propagate along conserved spatiotemporal trajectories.

## Discussion

We imaged mesoscopic spontaneous activity in the developing ferret visual cortex at high temporal resolution, revealing the dynamic properties of spontaneous network events. We found that the vast majority of events exhibited fast temporal dynamics, initiating in a local region and propagating across the imaged field of view. While some events were described well by a linear wave propagating through spatial modules, many were not. We observed a high degree of stereotypy in individual events, with more than half of events repeating multiple times throughout the recording. Furthermore, spatiotemporal activity patterns exhibited conserved trajectories over long timescales, with network events retaining information about past frames for as long as a second.

Our study builds on previous work that characterized the spatial correlations of the early neonatal cortex, but only hinted at the underlying temporal dynamics. In the visual cortex (Smith et al. 2018) as well as in other cortical areas early in development (Powell et al. 2024), spontaneous activity is typified by distributed, modular patterns, with correlated activity across millimeters. In the visual cortex, it was shown that the network correlation structure remained intact even after silencing feedforward connections, indicating that intracortical connections are sufficient to generate long-range correlations (Smith et al. 2018). This is supported by the finding that local excitation lateral inhibition models (LE/LI) are able to recapitulate the spatial characteristics of millimeter-scale correlation patterns using only local connections (Smith et al. 2018), consistent with the immaturity of long-range anatomical connectivity at this developmental stage (Durack and Katz 1996). However, these results are based on steady-state activity patterns of the model, thus their ability to describe more realistic conditions such as local spontaneous event initiation or spatially restricted input is unclear. In such conditions, the large-scale patterns observed both in the model and *in vivo* would necessitate the propagation of activity across the network. Our work uncovers previously unknown temporal dynamics of this activity by showing that the vast majority of spontaneous events *in vivo* have a dynamic component in which activity does in fact propagate out of the initial region of activation.

Inspired by the large body of work examining wave-like properties of spontaneous activity in the cortex, we attempted to describe the spontaneous events we observed using a linear wave model. Wave-like cortical activity has been observed in a number of studies including adult cats and monkeys (Muller et al. 2014; Sato, Nauhaus, and Carandini 2012; Nauhaus et al. 2012; 2009; Grinvald et al. 1994; Benucci, Frazor, and Carandini 2007).These results contrast with our finding that the majority of events were poorly described by a linear wave and exhibited more complex spatiotemporal dynamics, potentially reflecting differences in early development versus the mature cortex. Waves observed in the mature visual cortex have faster wavefronts than our observations (Benucci, Frazor, and Carandini 2007; Muller et al. 2014), thus likely relying on unmyelinated horizontal connections (Muller et al. 2018) which are yet immature in our ferrets (Durack and Katz 1996). Differences in recording modalities may also account for these results, as electrophysiological and voltage-sensitive dye measures of activity are sensitive to sub-threshold changes in activity, while calcium imaging is more sensitive to supra-threshold events. Wave-like activity has also been observed in local populations of neurons with 2-photon imaging in the mouse (Siegel et al. 2012) and the ferret (Smith et al. 2015), although given the differences in imaging field of view, direct comparison to our mesoscopic data is difficult. Interestingly, more complex non-linear waves, such as spiral waves, have been reported in cortex (Huang et al. 2010), although given the short duration and modular structure of the events we observe, an analysis of spiral structure in our dataset is likely to be inconclusive.

While the spatiotemporal sequences of spontaneous activity we observed were complex, the trajectories of propagating activity were strongly non-random. Approximately half of the 100 ms activity sequences we observed could be well described by a relatively small number (<8) of stereotyped sets of activity patterns that recurred across hours of spontaneous activity. Similarly structured event-event correlation matrices have been observed in *in vitro* networks of cultured neurons that exhibit avalanche criticality (Beggs and Plenz 2004). This suggests that there may be a common network-level model accounting for this behavior, but that remains to be explored in future work. Intriguingly, when we identified the most frequently observed spatial patterns, we found that activation of a particular spatial motif was predictable up to 100 ms prior to activation and the subsequent trajectory retained information about motif identity for nearly 1000 ms following activation. The tendency of cortical activity to converge on a relatively small number of states and retain state information suggests that the system is possibly in an attractor regime (Miller 2016). Attractor dynamics have been used to model mature visual cortex, including area MT (Pattadkal et al. 2024) and V1 (Priebe 2016), with our results suggesting they may also apply to early development at the modular scale.

In summary, our results reveal for the first time the complex spatiotemporal patterns of spontaneous activity in the early developing visual cortex. Even at this early stage of development, highly dynamic activity patterns are structured and non-random, with stereotyped spatial patterns that evolve in temporally predictable patterns. These structured sequences suggest mechanisms through which early spontaneous activity might support the development and refinement of mature modular cortical networks.

## Methods

### Experiments

#### Animals

All experimental procedures were approved by the University of Minnesota Institutional Animal Care and Use Committee and were performed in accordance with guidelines from the U.S. National Institutes of Health. We obtained six male and female ferret kits from Marshall Farms and housed them with jills on a 16 hour light/8 hour dark cycle. No statistical methods were used to predetermine sample sizes, but our sample sizes are similar to those reported in previous publications.

#### Viral injections

Viral injections were performed as previously described (Smith and Fitzpatrick 2016) and were consistent with prior work (Smith et al. 2018). We expressed GCaMP8m (Zhang et al. 2021) by microinjecting AAV1 expressing hSyn.jGCaMP8m.WPRE (University of Minnesota Viral Vector and Cloning Core) into layer 2/3 of primary visual cortex at P10-14, approximately 10-15 days before imaging. Anesthesia was induced with isoflurane (3-4%) and maintained with isoflurane (1-2%). Buprenorphine (0.01 mg/kg) and glycopyrrolate (0.01 mg/kg) were administered IM, as well as 1:1 lidocaine/bupivacaine SQ at the site of incision. Animal temperature was maintained at approximately 37°C with a water pump heat therapy pad (Adroit Medical HTP-1500, Parkland Scientific). Animals were mechanically ventilated and both heart rate and end-tidal CO2 were monitored throughout the surgery.

Using aseptic surgical technique, skin and muscle overlying visual cortex were retracted, and a small burr hole was made with a handheld drill (Fordom Electric Co.). Approximately 1 µl of virus contained in a pulled-glass pipette was pressure injected into the cortex at two depths (∼200 µm and 400 µm below the surface) over 20 min using a Nanoject-II (World Precision Instruments). The craniotomy was filled with 2% agarose and sealed with a thin sterile plastic film, affixed with Vetbond.

#### Cranial window surgery

On the day of experimental imaging, ferrets were anesthetized with 3–4% isoflurane. Atropine was administered SQ (0.2mg/kg). Animals were placed on a feedback-controlled heating pad to maintain an internal temperature of 37–38°C. Animals were intubated and ventilated, and isoflurane was delivered between 1% and 2% throughout the surgical procedure to maintain a surgical plane of anesthesia. An intraparietal catheter was placed to deliver fluids. EKG, end-tidal CO2, and internal temperature were continuously monitored during the procedure and subsequent imaging session. A 6–7 mm craniotomy was performed at the viral injection site and the dura retracted to reveal the cortex.

Five of the animals had a custom 3D printed insert that was placed directly onto the brain and craniotomy site, with one 4 mm cover glass (round, #1.5 thickness, Electron Microscopy Sciences) adhered to the bottom, and sealed to the skull using Kwik-Sil. This method was used to gently compress the underlying cortex and dampen biological motion during imaging. Alternatively, one animal used a custom titanium headplate that was affixed to the skull using Metabond and a titanium insert that was sealed with a snap-ring (5/16-in. internal retaining ring, McMaster-Carr), and a 4mm cover glass glued to the insert. Upon completion of the surgical procedure, isoflurane was gradually reduced (0.6–0.9%) and then vecuronium bromide (0.4 mg/kg/hr) mixed in an LRS 5% Dextrose solution was delivered IP to reduce motion and prevent spontaneous respiration.

#### Widefield epifluorescence imaging

Widefield epifluorescence imaging was performed with a sCMOS camera (Zyla 5.5, Andor) controlled by μManager (Edelstein et al. 2010). Images were acquired in global shutter configuration at 50 Hz with 4×4 binning to yield 640×540 pixels (Zyla) and additional offline 8x8 binning to yield 80x68 pixels. Spontaneous activity was captured in 10 to 20-min imaging sessions, with the animal sitting in a darkened room facing an LCD monitor displaying a black screen.

### Imaging preprocessing

#### Signal extraction for wide-field epifluorescence imaging

Motion correction was performed to correct for mild brain movement during imaging, which was done by registering each imaging frame to a reference frame. The ROI was manually drawn around the cortical region. The baseline F_0_ for each pixel was obtained by applying a rank-order filter to the raw fluorescence trace with a rank of 510 and a time window of 1425. The rank and time window were chosen such that the baseline faithfully followed the slow trend of the fluorescence activity. The baseline corrected spontaneous activity was calculated as

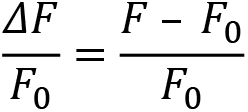

#### Deconvolution

We use a deconvolution method which we call Previous Frame Subtraction (PFS), an approach designed to remove calcium decay and highlights emerging activity. This is also referred to as numerical differentiation, as described in (Stern, Shea-Brown, and Witten 2020). We use the decay rate of GCaMP8m over 20 milliseconds, *γ*, as 89%. Then from every imaging frame, pixel-wise, we subtract the previous frame’s activity modulated by that decay.

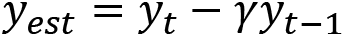

#### Event segmentation

We first determined active pixels on each frame by finding a threshold for each pixel that is 4 s.d. above its mean value. To remove spuriously active pixels, active pixels were required to be part of a contiguous area of at least 0.028mm^2^, using morphological operations of binary erosion followed by binary dilation on the thresholded data. An event was then defined as any contiguously active time period, with an inactive frame (defined as a frame without any active pixels) on either side. Finally, we only analyzed events that activated at least 1mm^2^ of cortex across its entire duration, in order to focus on large spontaneous evens with multiple active modules, as in prior work (Smith et al. 2018).

### Event metrics

#### Active area

The sum of all pixels that were active during the event at least once, measured in mm^2^.

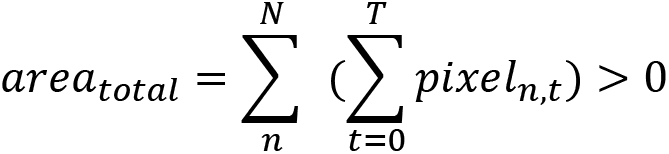

#### Propagation area (PA)

The area of additional activity after the onset frame. An exclusion radius of 400μm from onset activity was applied, and the total area is as above.

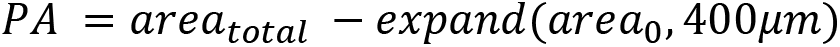

#### Linear wavefront fitting

A linearly traveling wavefront was computed by projecting the activity of an event onto a linear traveling wave with variables velocity, direction, and starting time, similar to (Smith et al. 2018). A pixel was considered activated at time *t* if the traveling wave passed it during the discrete time step *t*, and only the onset time was considered. The variables were estimated by minimizing the mean squared error (MSE) between the projected, continuous *t* values and observed *t* values, using an initial brute force search and then fine-tuning the values using an optimization algorithm.

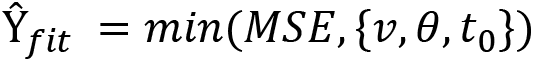

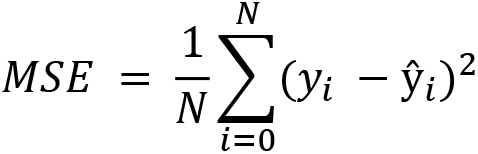

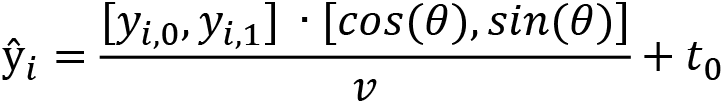

To determine if an event had a linear fit significantly better than chance, we permuted the order of frames for each event and fit the permutations. We found the fraction of permutations the observed MSE value performed better than, referred to as the “waviness index” in (Siegel et al. 2012) and considered anything above 95% as significant. To ensure sufficient numbers of possible permutations for accurate comparison, we restricted our analysis to events with 5 or more frames.

To determine the maximum velocity we could detect with our imaging parameters, we considered our imaging framerate of 50Hz and the size of each animal’s imaging window. Given a 3mm imaging window, activity could be observed to shift over the course of one frame (20 ms) with a maximum velocity of ∼150mm/s. Thus, we excluded any events with fit velocities greater than this maximum value.

The estimated propagation directions in our data appeared to have a symmetric bimodal distribution where the activity traveled across the imaging FOV roughly along an axis in either direction. To test this, we converted estimated propagation angles into axial space.

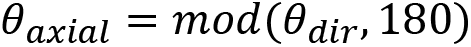

Then we applied the Rayleigh test for significant axial bias. If an animal had a significant axis of propagation, we further tested if there was a symmetric but uneven direction preference by comparing the number of linear events propagating along either direction (using +-90 degree bins around the mean axis) and performing a binomial significance test.

#### Event repetitions and clustering

To identify repeated spatiotemporal patterns (Fig 4), we identified a subset of events with the same number of datapoints (five frames). Unlike previous analyses that used thresholded activity, here we used deconvolved data, Gaussian bandpass filtered (s.d.: *s*low = 94 µm, *s*high = 560 µm), in order to emphasize the modular patterns and mitigate noise. We computed the event-event correlation matrix using Pearson correlation coefficient. We determined if an event was repeated with a permutation test. Cortical activity has a consistent rising and falling pattern for each event, so we controlled for this by permuting within a time step, i.e., all of the first frames were permuted. Thus, we took 1000 within-time step permutations, computed 1000 correlation matrices, and the 99th percentile of the permuted correlations was used as a threshold for a significant correlation. We applied this threshold to determine how many repetitions each event had. With these repeated events, we utilized a greedy algorithm to find clusters of patterns. That is, the event with the highest number of repeats defined the first group as this set of repeated events. These were excluded on the next iteration and then the second group was found, etc. Across animals, clusters were included in the following analyses if they included at least 10 events. The fractional range of each cluster (Fig. 4H) was computed as

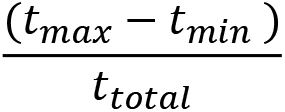

The fraction of bins (Fig. 4H) was computed as

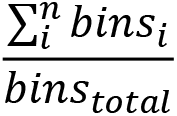

where *bins_i_* is a ten minute segment of the imaging session that had at least one instance of the cluster.

#### Template frame selection and group membership

To identify spatial activity motifs (Fig. 5), we aggregated the frames of all events that were 200 ms or shorter. In order to target modular activity and remove noise, each frame was fit to a two-dimensional multi-Gaussian function, giving us a set of “template patterns”. We computed a threshold for each template pattern by taking correlation values between the template and translated versions of the template, shifted in xy space up to one standard deviation away from the center of the Gaussian fit, and used the lowest correlation value of these translations as the threshold.

We then computed the correlation values between each frame of activity and all template patterns. Using a greedy algorithm (similar to *Event repetitions and clustering*), we took the template pattern with the most correlated frames, and used this to identify events containing those frames, which define the first group. We removed those events and repeated the procedure to identify the second group, etc. The greedy method is restricted to only using one frame per event, thus, if two frames were matched in a single event, we took the highest correlated frame to the template to determine group identity. In order to compare across animals, we used the top seven groups for all animals. This cutoff was chosen to ensure that every group across animals had at least 15 repeats per group, and such that every animal used the same number of groups for across-animal comparisons.

#### Classification accuracy for template groups

We independently decomposed each time point into its first eight principal components. Groups were matched to have the same number of samples through subsampling, and we subsampled 100 times. To find the reported classification accuracy from −2000ms to 2000ms, we took each matched and subsampled set and computed the mean of its five-fold cross-validation test accuracy. This yielded a distribution of classification accuracies for all 100 subsamples.

We used a shuffled dataset to control for any unknown biases in classification accuracy. In order to do this, we took each of the 100 subsamples and permuted group labels for each one 100 times. Each subsample was compared against the 100 permutations of that subsample, and was considered significant if classifier performance was greater than the 95th percentile of the permuted data. Finally, a time point was considered significantly classified if 95% of subsamples were significantly classified.

## Author Contributions

Conceptualization: LK, GBS Methodology: LK, AS, GBS Investigation: LK

Formal Analysis: LK

Visualization: LK Funding acquisition: GBS Supervision: AS, GBS

Writing – original draft: LK

Writing – review & editing: LK, AS, GBS

## Acknowledgments

The authors wish to thank Nic Glewwe, Matt Paruzynski, Jack Kapler, and Sophie Bowman for surgical assistance, Ryan Holland for coding support, Deano Farinella and Harishankar Jayakumar for their optical expertise, and members of the Smith and Sederberg labs for helpful discussions. All viral vectors used in this study were generated by the University of Minnesota Viral Vector and Cloning Core (Minneapolis, MN).

This work was supported by these funding sources: National Eye Institute (R01EY030893) (GBS) Whitehall Foundation (2018-05-57) (GBS) National Science Foundation (IIS-2011542) (GBS)

NIH Training Program in Translational Vision Sciences T32EY025187 (LK) NIMH Training Program in Computational Neuroscience T32MH115866 (LK)

---

**Supplemental Figure 1.**
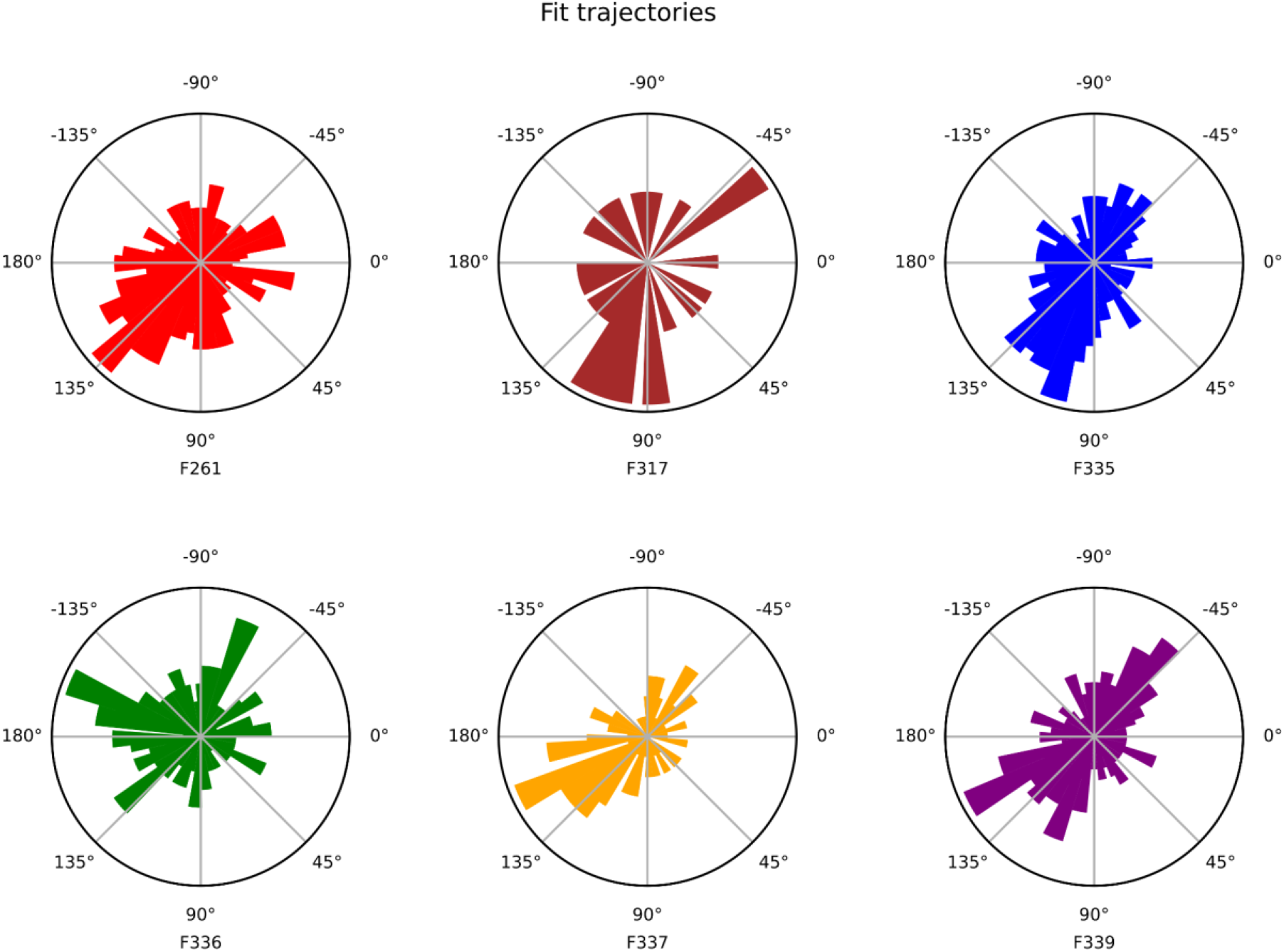
Expanded data from Fig. 3. Each subplot is data from one animal and shows the distribution of fit trajectories for all events that were significantly a linear wave. Animals are color-coded for easier referencing back to the main figure.

**Supplemental Figure 2.**
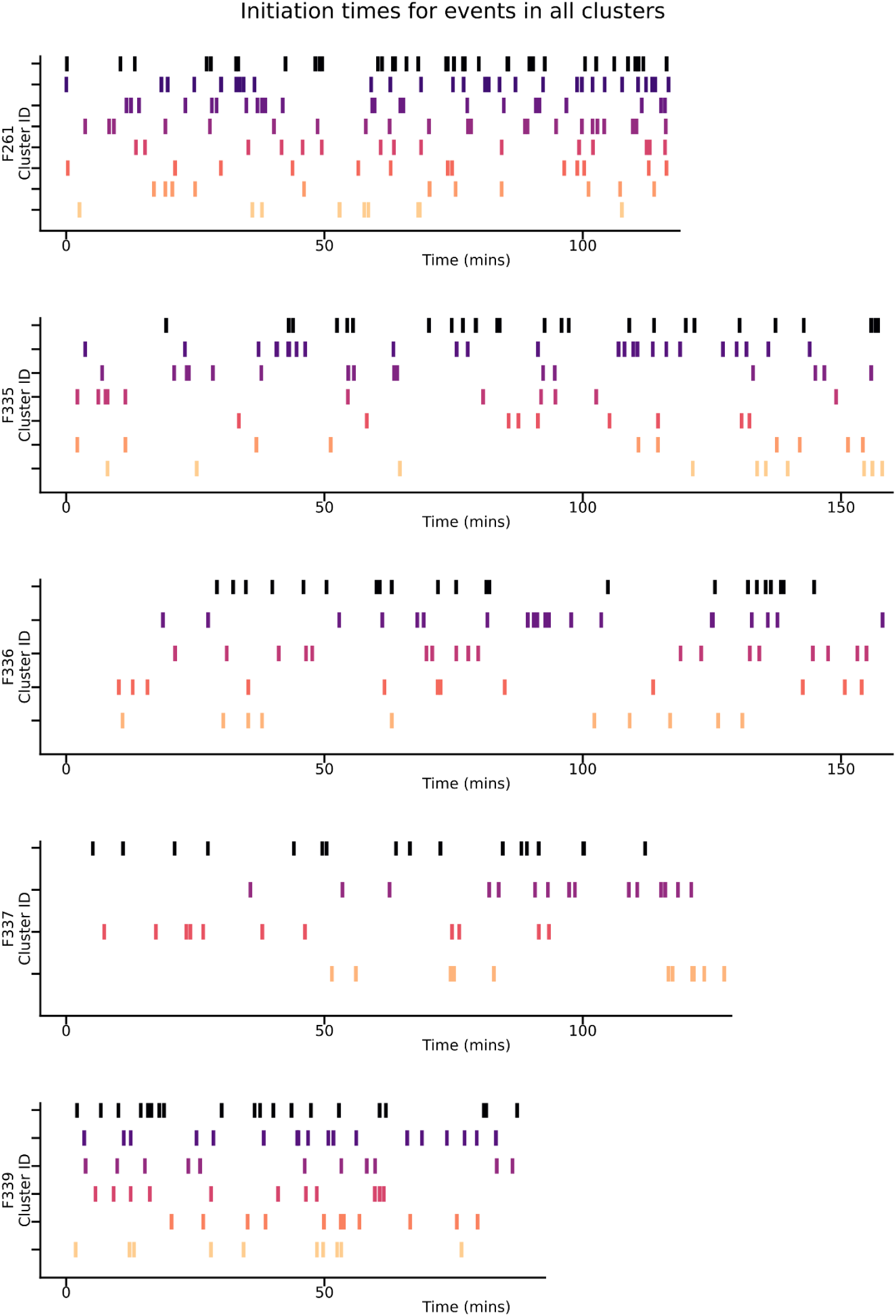
Figure 4G expanded to all animals. Each plot is one animal and their imaging session. The initiation times are plotted for events in each cluster, where the cluster is distinguished by row and color.

**Supplemental Table 1.**

| Ferret | Injection age | Imaging age | Expression time (days) | FOV area (mm<sup>2</sup>) |
| --- | --- | --- | --- | --- |
| F261 | 14 | 24 | 10 | 5.5 |
| F317 | 12 | 23 | 11 | 5 |
| F335 | 11 | 23 | 12 | 6 |
| F336 | 11 | 25 | 14 | 6.5 |
| F337 | 11 | 28 | 17 | 8.9 |
| F339 | 11 | 25 | 14 | 5.8 |

**Supplemental Table 2.**

| Ferret | Total events | Significant linear wave | Fraction significant linear wave | Axial (180) Rayleigh test p-value | Binomial test p-value |
| --- | --- | --- | --- | --- | --- |
| 261 | 867 | 462 | 0.53287 | 0.0001167 | 3.66781E-05 |
| 317 | 100 | 30 | 0.3 | 0.1615 | - |
| 335 | 934 | 396 | 0.42398 | 8.043E-10 | 0.000370051 |
| 336 | 627 | 161 | 0.25678 | 0.4107 | - |
| 337 | 441 | 102 | 0.23129 | 1.089E-07 | 0.000996403 |
| 339 | 824 | 281 | 0.34102 | 6.248E-10 | 0.360233 |
| All | 3793 | 1432 | 0.37754 | - | - |

